# High-resolution transcriptome atlas and improved genome assembly of common buckwheat, *Fagopyrum esculentum*

**DOI:** 10.1101/2020.11.14.382903

**Authors:** Aleksey A. Penin, Artem S. Kasianov, Anna V. Klepikova, Ilya V. Kirov, Evgeny S. Gerasimov, Aleksey N. Fesenko, Maria D. Logacheva

## Abstract

Common buckwheat (*Fagopyrum esculentum*) is an important non-cereal grain crop and a prospective component of functional food. Despite this, the genomic resources for this species and for the whole family Polygonaceae, to which it belongs, are scarce. Here, we report the assembly of the buckwheat genome using long-read technology and a high-resolution expression atlas including 46 organs and developmental stages. We found that the buckwheat genome has an extremely high content of transposable elements, including several classes of recently (0.5-1 Mya) multiplied TEs (“transposon burst”) and gradually accumulated TEs. The difference in TE content is a major factor contributing to the 3-fold increase in the genome size of *F. esculentum* compared with its sister species *F. tataricum*. Moreover, we detected the differences in TE content between the wild ancestral subspecies *F. esculentum* ssp. *ancestrale* and buckwheat cultivars, suggesting that TE activity accompanied buckwheat domestication. Expression profiling allowed us to test a hypothesis about the genetic control of petaloidy in buckwheat. We showed that it is not mediated by B-class gene activity, in contrast to the prediction from the ABC model. Based on a survey of expression profiles and phylogenetic analysis, we identified the MYB family transcription factor gene tr_18111 as a potential candidate for the determination of conical cells in buckwheat petaloid tepals. The information on expression patterns has been integrated into the publicly available database TraVA: http://travadb.org/browse/Species=Fesc/. The improved genome assembly and transcriptomic resources will enable research on buckwheat, including practical applications.

## Introduction

Common buckwheat (*Fagopyrum esculentum*) is an important non-cereal grain crop of particular importance in Russia, China and Ukraine, with production of ~ 2 million tons. With the growing interest in healthy lifestyles, buckwheat has received a great deal of attention worldwide as a component of functional food in recent years (for review, see (Giménez-Bastida and Zieliński, 2015; Alvarez-Jubete *et al.*, 2010)). Buckwheat is highly resistant to soil contamination by aluminium and lead and is a promising phytoremediator (Ma *et al.*, 1997; Honda *et al.*, 2007). Its productivity as an agricultural plant is, however, limited by several unfavourable traits: obligate insect outcrossing pollination, an extended period of flowering and susceptibility to drought and freezing. Novel methods of breeding that involve marker-assisted and genomic selection have the potential to overcome these limitations. For this purpose, genomic information is highly desired. A draft assembly of the buckwheat genome was reported in 2016 (Yasui *et al.*, 2016); however, it is based only on Illumina technology and shows moderate continuity (contig N50 1 Kb, scaffold N50 = 25 Kb). This assembly is likely to encompass most coding regions and is suitable for the analysis of single genes (though with some limitations, see, e.g., Lei *et al.*, 2017). Many researchers opt not to use this assembly as a reference for transcriptomic studies on buckwheat and instead generate their own de novo transcriptome assemblies (Xu *et al.*, 2017; X., Fang *et al.*, 2019; Z., Fang *et al.*, 2019). This situation decreases the accuracy of gene expression estimates and hampers the comparison and meta-analysis of the results. Thus, the generation of a more contiguous assembly for the buckwheat genome complemented with transcriptomic information is urgently needed (Joshi *et al.*, 2019).

Sequencing technologies have been responsible for tremendous progress; one of the key innovations of these technologies in recent years was the development of NGS platforms capable of generating long reads. This is of special importance for plant scientists because plant genomes are large and complex and are shaped by multiple segmental and whole-genome duplications and transposable element (TE) activity. Thus, they cannot be reliably assembled using 2^nd^-generation NGS technologies, which generate reads with a maximum length of 100-300 bp. The utility of long-read technology for the assembly of complex genomes has been demonstrated in several plant species (Schmidt *et al.*, 2017; Murigneux *et al.*, 2020). Long-read technologies have encouraged scientists to conduct studies that were impossible even with the gold-standard quality genome of *Arabidopsis thaliana* due to the complex rearrangements caused by transposition. On this basis, we performed the sequencing, de novo assembly and annotation of the buckwheat genome using 3^rd^-generation technologies – SMRT and nanopore sequencing – and generated a transcriptome map of *F. esculentum* from 46 organs and developmental stages. We demonstrate the utility of detailed transcriptome profiling for testing and generating hypotheses about the involvement of genes in specific processes, particularly the genetic basis of floral organ identity and anthocyanin synthesis. We show that gene expression patterns are a valuable addition to sequence-based information.

We expect that this newly generated genome and transcriptome resource will be useful for the development of buckwheat genetics and breeding. Buckwheat belongs to an isolated group of flowering plants, the non-core Caryophyllales, which are distantly related to model species (Yao *et al.*, 2019). While their sister clade, the core Caryophyllales, contains several plants with well-characterized genomes (sugarbeet (Dohm *et al.*, 2013) and quinoa (Jarvis *et al.*, 2017)), genomic and transcriptomic resources for the non-core Caryophyllales are lacking. Thus, the development of a high-resolution transcriptome map will provide a resource for comparative transcriptomic analyses in the phylogenetic context.

## Results and discussion

### Assembly and annotation

The Dasha cultivar was selected for reference genome assembly. Dasha is a recently developed fast-growing determinate cultivar characterized by resistance to lodging and a high photosynthesis efficiency (Fesenko *et al.*, 2018). The assembly process consisted of three stages: contig assembly, scaffolding and gap closing. For contig assembly, a Newbler assembler was used. This assembler, initially developed for 454 data, also performs well for other types of sequencing data, including merged Illumina paired 250-bp or 300-bp reads. The 250-bp Illumina reads and mate-pair reads provided a backbone for the assembly; further improvement of the assembly, including gap closing, was performed using long reads generated by SMRT CCS technology on the Pacific Bioscience platform (Table S1). The final assembly consisted of 88 078 scaffolds with a total length of 1.2 Gb and N50 ~ 180 Kb (Table S2). The N50 of this assembly represents a 39x improvement compared with that of the previous assembly at the level of contigs. A search of the conserved gene set (BUSCOs) also showed high continuity of the assembly: the percentage of complete genes was 98.6%, and those of fragmented and missing genes were both 0.7%. For comparison, these metrics for the previous assembly were 84.2%, 11.5 and 4.3% (Table S3). The annotation yielded 29 514 genes; the annotation metrics are summarized in Table S4. The majority of the genes (97.5%) were supported by RNA-seq (Table S4 and text below). The parameters of the coding fraction of the genes (CDS length, exon length) of buckwheat were similar to those of other plants, while the introns of buckwheat were longer, which is in accord with its larger genome size (Figure 1a). The number of genes was close to that found in sugar beet (Dohm *et al.*, 2013), which belongs to Caryophyllales along with buckwheat, and in *A. thaliana* and was generally within the range typical for the genomes of plants that have not undergone a recent whole-genome duplication. A total of 24 765 buckwheat genes were classified into 12 627 orthogroups. Notably, the buckwheat genome contains a smaller number of orthogroups that include one gene and a larger number of orthogroups that include two or more genes (Figure 1b). This pattern is not, however, compatible with that observed in tetraploids (see, e.g., Leushkin *et al.*, 2013); taking into account the unusually high content and activity of TEs in the buckwheat genome (see below), we conclude that it is a result of segmental duplications caused by TEs. The distribution of gene ontology categories was similar between buckwheat and *Arabidopsis* (except for the “secondary metabolic process” category, which includes more genes in *Arabidopsis*, presumably due to its ability to synthesize glucosinolates) (Figure 1c). Additionally, a search for GO enrichment in the genes belonging to the 1-to-many orthogroups (one gene in *Arabidopsis* and two or more in buckwheat) did not reveal any enrichment (FDR < 0.01). This shows that duplications are not confined to specific gene groups. A total of 1 425 buckwheat genes were classified as transcription factors (TFs, Table S5); the total number and distribution of TF classes were similar to those of other caryophyllids (1 058 in *Beta vulgaris*, 1 259 in *Amaranthus hypochondriacus*) and *A. thaliana* (1 717). The extranuclear genomes – plastid and mitochondrial – were assembled using different procedures and are reported elsewhere (Logacheva *et al.*, 2020).

**Figure 1.**
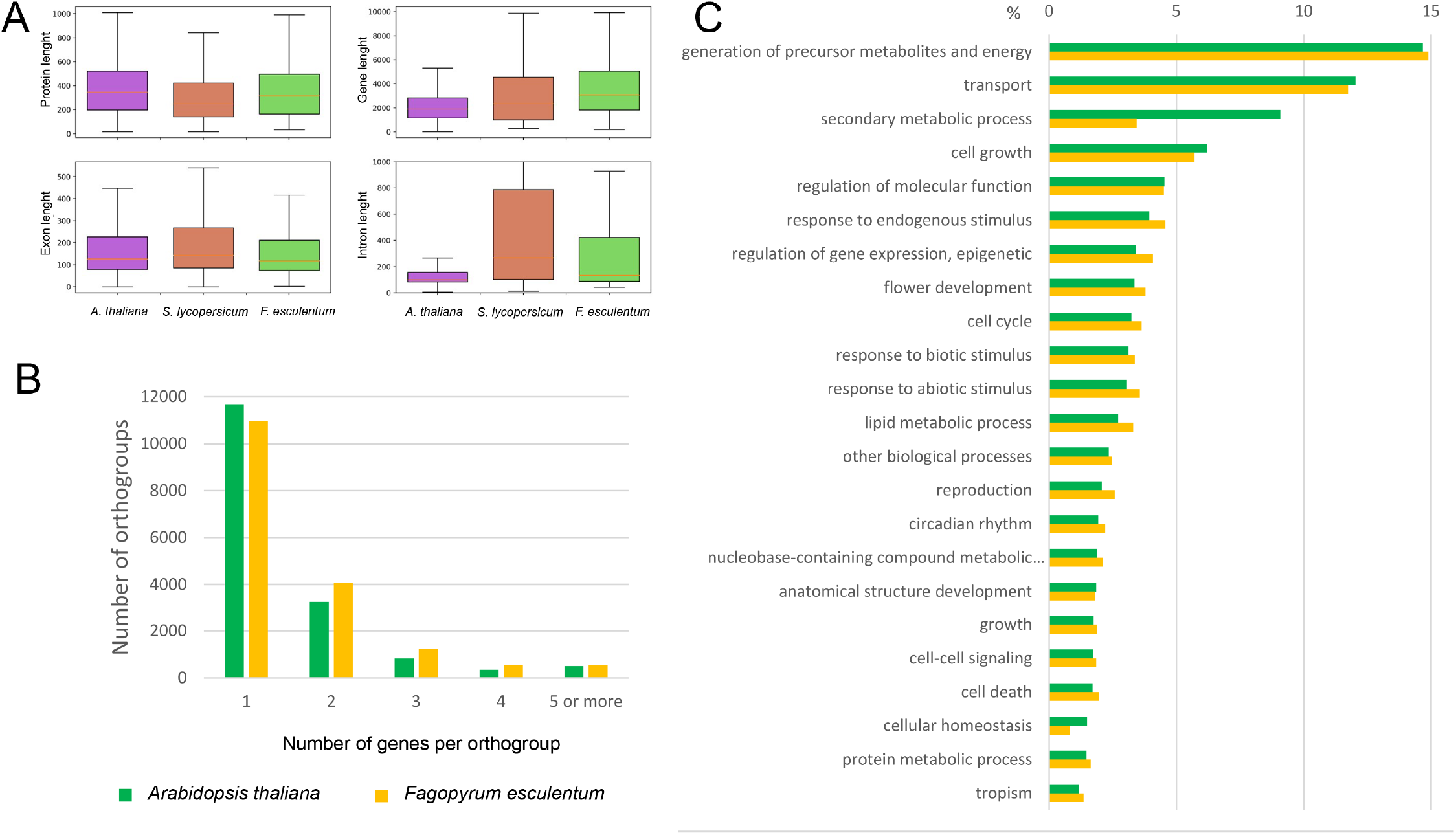
a) Annotation of *F. esculentum* compared with those of *A. thaliana* and *S. lycopersicum* in terms of the gene, exon and intron length distribution b) The number of genes per orthogroup in *Fagopyrum esculentum* and *Arabidopsis thaliana* c) The distribution of GO categories (biological function) *in Fagopyrum esculentum* and *Arabidopsis thaliana*.

### Transposable elements and the evolution of genome size in the genus Fagopyrum

The genus *Fagopyrum* includes approximately 25 species, which fall into two clades: cymosum (large achene) and urophyllum (small achene) (reviewed in (Ohsako and Li, 2020)). The genome sizes of the two clades vary significantly despite showing the same chromosome number in most species (Nagano *et al.*, 2000); the basis of this variation is unknown. *F. esculentum* is a member of the large-seeded clade; its genome is ~3 times larger (1.39 pg/1C) than that of the closely related species *F. tataricum* (0.56 pg/1C), while the chromosome number is the same in these species (Neethirajan *et al.*, 2011). Polyploidy and TEs are the main drivers of plant genome size variability across species (Bennett and Leitch, 2005; Vitte and Panaud, 2005). Because both species are diploids and they show the same chromosome number (2n=2x=16), we hypothesized that TE activity may be responsible for the increase in *F. esculentum* genome size. To obtain preliminary insight into the genome-wide differences in the repeatome composition between the two species, we performed the comparative clustering of genomic NGS reads followed by cluster annotation using RepeatExplorer software (Novak *et al.*, 2013; Novák *et al.*, 2017). This analysis revealed a similar genome abundance of satellite (3.45% and 4%) and class II (DNA transposons, 1.97% and 1.98%) repeats, while the portion of the genome occupied by retrotransposons (RTEs, class I repeats) was ~2 times greater in *F. esculentum* (64.8%) than in *F. tataricum* (30.7%) (Figure 2a). Then, we took advantage of the current *F. esculentum* genome sequence assembly to obtain deeper insight into LTR RTE diversity and compared these data with those of *F. tataricum* (http://mbkbase.org/Pinku1/FtChromosomeV2.fasta.gz, (Zhang *et al.*, 2017). The de novo prediction of full-length LTR retrotransposons in the two genome assemblies revealed 14 435 and 4 891 sequences for *F. esculentum* and *F. tataricum*, respectively, showing good concordance with the NGS-based estimation of TE abundance differences. The further classification of RTEs indicated that ~70% of all RTEs belonged to the Ty3/Gypsy superfamily. The family-based classification of all RTEs revealed striking differences in the copy numbers of distinct Ty3/Gypsy and Ty1/Copia families between the two species (Figure 2b, c). Athila Ty3/Gypsy was the most prevalent (42%) family in the *F. esculentum* genome, including ~25 times more RTEs (6117) than the number found in *F. tataricum* (241) (Figure 2b). In contrast, 59% of *F. tataricum* RTEs (2911) belong to the Tekay Ty3/gypsy RTEs (Figure 2b), although the number of identified RTEs of this family was almost identical between the two species. Similarly, Ty1/Copia RTEs showed discrepancies in the copy numbers of distinct families between the genomes of two species, with the Ale, SIRE, TAR and Ikeros families being the most prevalent in *F. esculentum*, while the Tork family is the largest family in *F*. *tataricum,* although the copy number is almost the same between the two genomes. This suggests that the RTEs of several families have accumulated at a higher rate in *F. esculentum* than in *F. tataricum*. We then asked whether the insertion time of RTE activity differed between these species. The obtained estimates clearly showed significant differences (Fisher’s exact test for count data, p-value < 2.2e-16) between the species, with 26.6% (3 834 complete RTEs) of the RTEs of the *F. esculentum* genome having been inserted during the last 0.5 million years (‘recent’ insertions), while only 9.6% of *F. tataricum* RTE insertions were classified as ‘recent’ insertions. Surprisingly, the comparison of insertion times for different RTE families revealed that Athila accumulation was characterized by gradual dynamics in both species, without a burst of recent activity in *F. esculentum* (Figure 2e). A similar situation was observed for the CRM and TAR families. In contrast, the Ale, Ikeros, and Tork families and the SIRE family showed signatures of bursts of recent activity in *F. esculentum* and *F. tataricum*, respectively. Thus, the greater *F. esculentum* genome size compared with that of *F. tataricum* is influenced by the generally higher rate of transposition activity and gradual accumulation of Athila members, together with the burst of recent activity of Ale, Ikeros and Tork family members. Along with the comparison of *F. tataricum* and *F. esculentum*, which allowed us to reveal the basis of the rapid genome size change, we surveyed the TE content within *F. esculentum*. We analysed WGS data for 10 cultivars and the ancestral subspecies *F. esculentum* ssp. *ancestrale* and found that the abundance of TEs differed between the cultivars and *F. esculentum* ssp. *ancestrale* and between the cultivars. (Figure S1). These differences were correlated with the breeding history of the cultivars; for example, the Kazanka (Kaz) and Dialog (Dia) cultivars, which were clustered together in the TE abundance tree (Figure S2), are indeed closely related, with Kazanka being one of the progenitors of Dialog (Fesenko, 2009).

**Figure 2.**
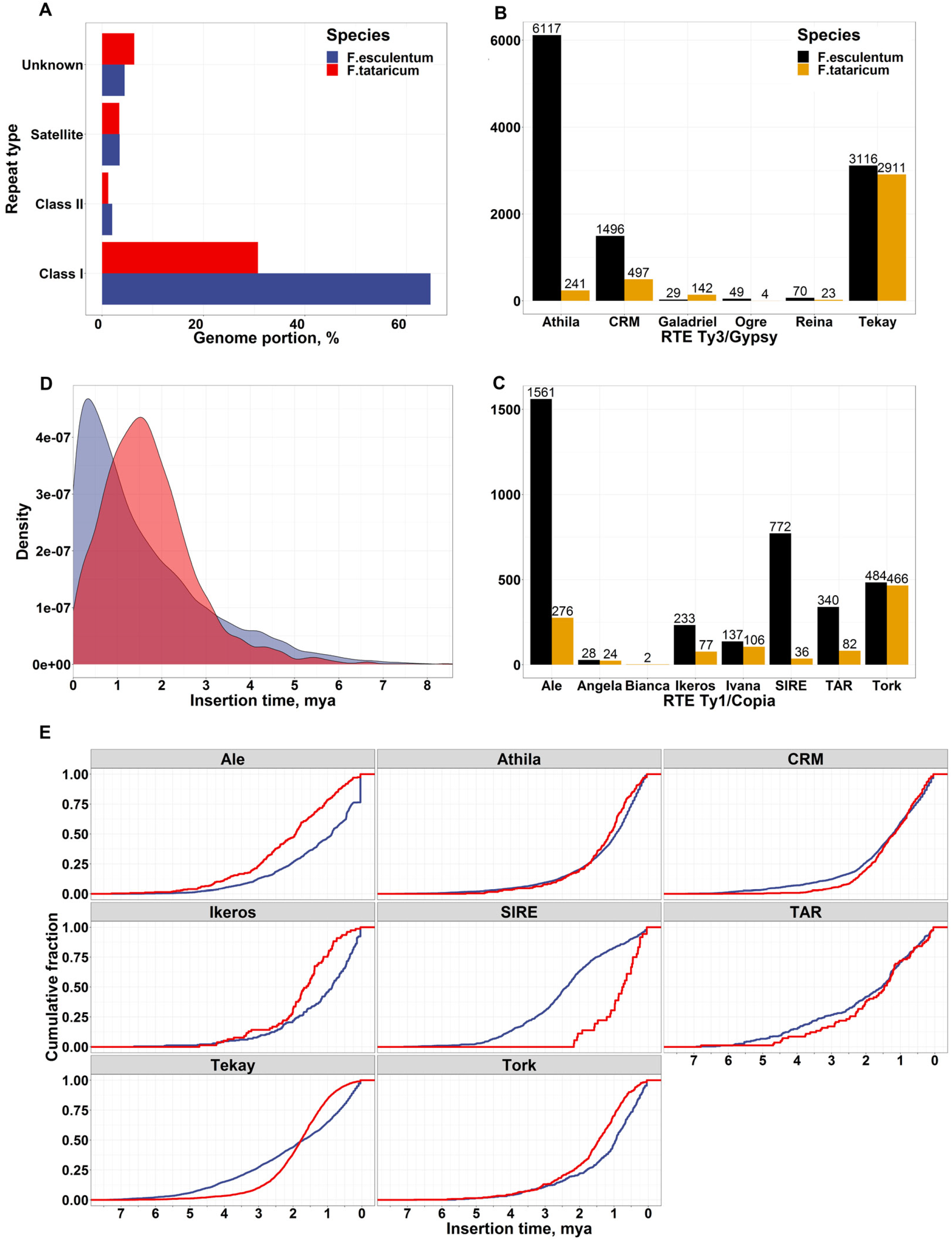
a) Comparison of genome abundance of different types of repeats in the *F. esculentum* and *F. tataricum* genomes based on the RepeatExplorer (Novak *et al*., 2013) clustering of NGS genomic reads. b) and c) Number of full-length Ty3/Gypsy and Ty1/Copia LTR RTEs of different families based on prediction in the assembled genome sequences. d) Insertion time comparison for *F. esculentum* (blue) and *F. tataricum* (red) LTR RTEs. e) Cumulative plot of the insertion time for RTE families.

### Buckwheat transcriptome atlas and FescTraVA database

Gene expression information is integral for many areas of plant biology; in particular, it helps to reveal the role of genes in growth, development and stress response (see e.g. (Zhao *et al.*, 2014), (Shumayla *et al.*, 2017)), to find the loci associated with agriculturally important traits (Galpaz *et al.*, 2018), to infer phylogenetic relationships between species (Guo *et al.*, 2020). With this premise we constructed transcriptome map of buckwheat based on newly generated genome sequence and annotation. We selected 46 samples from different organs and developmental stages of buckwheat, ranging from seeds and seedlings to senescent organs (Table S6). Since buckwheat exhibits two forms of flowers – pin (long style and short stamens) and thrum (short style and long stamens) – most samples of flowers and floral parts were collected separately for the two forms. The transcriptome is highly dynamic, especially in plants, in which it depends greatly on environmental conditions and the circadian cycle. Taking this into account, the growing and collection of samples for the transcriptome map was performed under the same conditions and at the same time of the day. For each sample, > 20 million reads were obtained, and the average number of reads per sample was 12 million. The correlation coefficient of the samples lies in the range 0.85-1.00 with an average of 0.96 (Table S7). The clustering of the samples reflects their biological similarity (Figure 3a). The number of genes expressed in each sample was similar, and the greatest numbers were observed in seeds and flowers (Figure 3b). The number of genes expressed in at least one sample was 28 770 (97% of annotated genes), while 12 172 genes were expressed in all samples, and 744 genes were not expressed in any (Figure 3c). The general patterns of gene expression were similar to those observed in *Arabidopsis* (Klepikova *et al.*, 2016). The estimation of gene expression breadth using Shannon entropy showed that most genes exhibited broad expression patterns (Figure S2). The genes in the fraction with the broadest expression patterns were enriched in the GO categories “photoperiodism, flowering” and “protein transporter activity”, while those in the fraction with the narrowest expression patterns were enriched in “serine-type endopeptidase inhibitor activity”, “negative regulation of endopeptidase activity”, and “endomembrane system” (Table S8).

**Figure 3.**
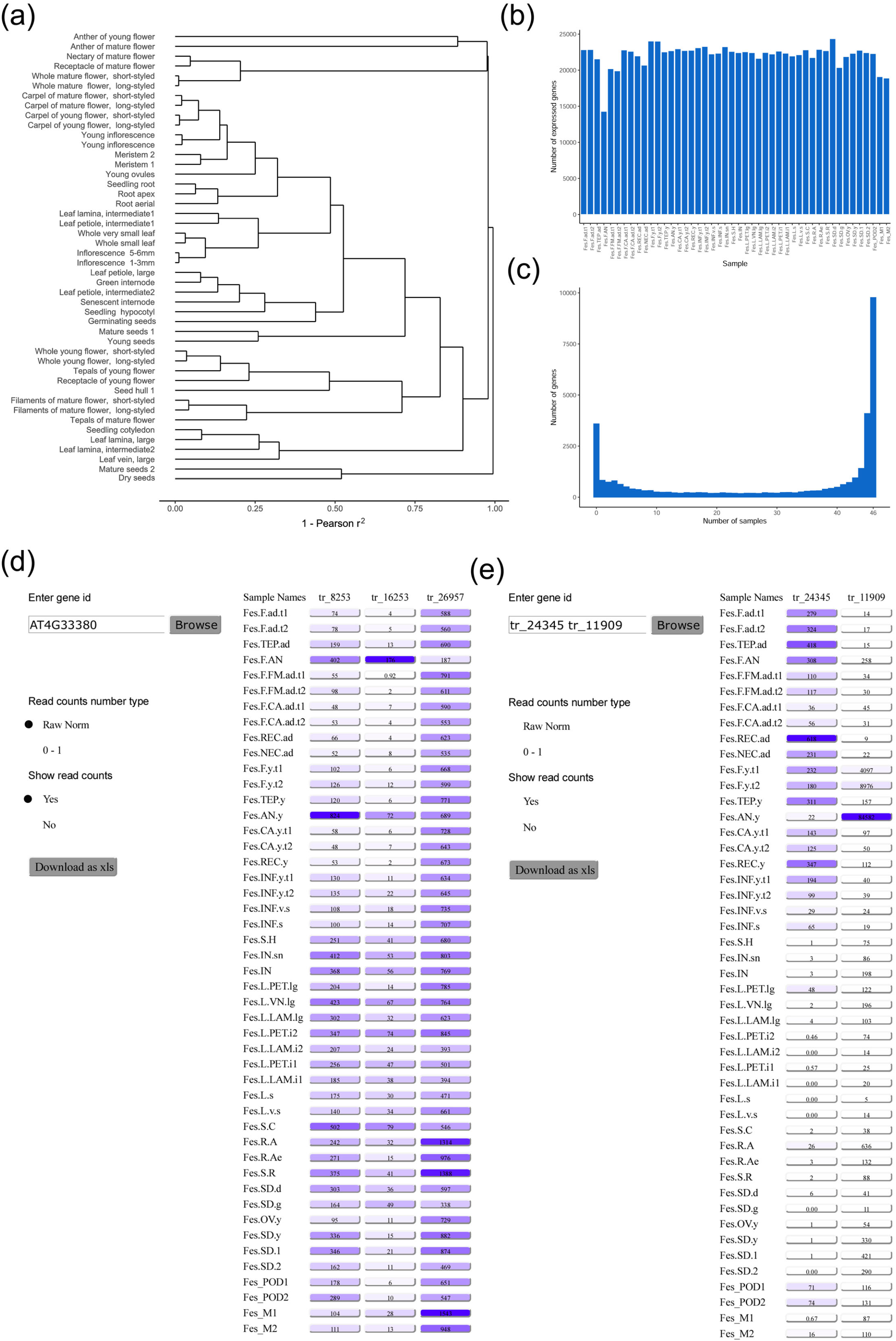
a) Clustering of the samples (the distance measure is 1 - square of Pearson correlation coefficient, the expression values for replicates are averaged). b) The number of genes expressed in each sample. c) The specificity of gene expression. d) The interface of the FescTraVA database, showing an example of a search with an *Arabidopsis* TAIR gene ID e) an example of a search with a buckwheat gene ID. Numbers in boxes denote normalized read counts.

RNA-seq data, especially high-resolution transcriptome maps, are an important resource for the identification of stably expressed genes, which are useful as reference genes for qRT-PCR (Zhou *et al.*, 2017; Machado *et al.*, 2020). We used the standard deviation of gene expression divided by mean expression (standard deviation; SD)/mean) as a measure of expression stability, and we identified 20 transcripts with SD/mean ratios < 0.2, 205 with ratios < 0.25, and 720 with ratios < 0.3. In a previous study, several gene orthologues of stably expressed *Arabidopsis* genes (Czechowski *et al.*, 2005) were evaluated in five organs of buckwheat using qRT-PCR (Demidenko *et al.*, 2011). Ten transcripts corresponded to these genes, and only one of them – tr_15881 (an orthologue of AT4G34270) – was stably expressed. The organs where expression showed the greatest differences were the anthers and roots (Figure S3), which were not sampled in the abovementioned study. This stresses the importance of balanced and high-resolution transcriptome maps for the assessment of gene stability.

To make the information on gene expression levels readily available to the research community, we summarized the results in the TraVA database: http://travadb.org/browse/Species=Fesc/. This database uses the same intuitive user-friendly interface as the transcriptome atlas of *A. thaliana* published earlier (Klepikova *et al.*, 2016). For convenience, the database can be searched according to either the identifier of the buckwheat transcript or the identifier (common name or AT*G* TAIR identifier) of the homologous *A. thaliana* gene. When the *A. thaliana* gene is a member of an orthogroup including several buckwheat genes, the expression profiles for all of these genes are shown (Figure 3d, e). The data can be downloaded in PNG and XLS formats.

### The potential of comparative transcriptomics for the inference of gene function

High-resolution gene expression maps are a useful tool for the generation and testing of hypotheses about gene function in non-model organisms. A commonly used approach is the identification of an orthologue of a gene of interest in a model organism, where the functions of the orthologues are assumed to be similar (the so-called orthologue conjecture) (Gabaldón and Koonin, 2013; Altenhoff *et al.*, 2012). Thus, the efforts of comparative genomics are focused on the identification of orthologues (Tulpan and Leger, 2017). However, this approach has several limitations. The most critical aspect of plant science is the complexity of plant gene families, which are shaped by multiple whole-genome and segmental duplications. This leads to the impossibility of inferring 1-to-1 orthologues and makes it necessary to work with orthogroups – the sets of genes descended from a single gene in the last common ancestor. The diversification of gene function within an orthogroup is a common way in which new structures and pathways emerge in plants (e.g., Yamaguchi *et al.*, 2006; Ober, 2005). However, based on sequence similarity, it is usually impossible to identify genes that share the same function in two species because subtle differences at the coding sequence level can contribute greatly to changes in function (a single amino acid change can turn a repressor to an activator) (Hanzawa *et al.*, 2005).

The transcriptome atlas enables the comparison of the expression profiles of the genes within an orthogroup, thus allowing the prioritization of the hypotheses regarding their functional correspondence. In a previous study, we constructed transcriptome atlases for *Arabidopsis* (Klepikova *et al.*, 2016) and tomato (Penin *et al.*, 2019) using the same approaches and a similar set of samples employed for buckwheat; these species represent two major groups of eudicots (rosids and asterids) and provide a basis for the comparative analysis of gene expression profiles. As an illustration of the novel information that the atlas can provide, we will focus on at the orthogroup including the *APETALA1* gene. The buckwheat genome carries three orthologues of the *AP1* gene. They share high similarity (74-77% at the nucleotide sequence level) and likely arose via whole-genome or segmental duplication after the diversification of core and non-core Caryophyllales (Figure 4a). The comparison of their expression profiles showed that the expression of all of these genes was confined to the flower; however, the patterns and levels of expression were different (Figure 4a). tr_22431 showed the highest expression level in tepals (the structures surrounding the reproductive organs of the flower in buckwheat, which are similar to petals and sepals; see discussion below), while two others showed the highest expression in the anthers and the receptacle of the developing flower. In *Arabidopsis*, *AP1* is known to regulate sepal and petal development (Irish and Sussex, 1990, p.1). Based on the expression profiles, the most plausible candidate for this function was tr_22431. Indeed, a study by Liu *et al.* (2019), who performed a complementation assay with an *Arabidopsis ap1* mutant and a buckwheat *AP1* orthologue (corresponding to tr_8095 from our annotation, 100% identity), showed a limited conservation of function between *Arabidopsis AP1* and this buckwheat gene. This shows the congruence of functional studies and expression profiles and calls for the widespread application of the buckwheat transcriptome atlas for functional genomics.

**Figure 4.**
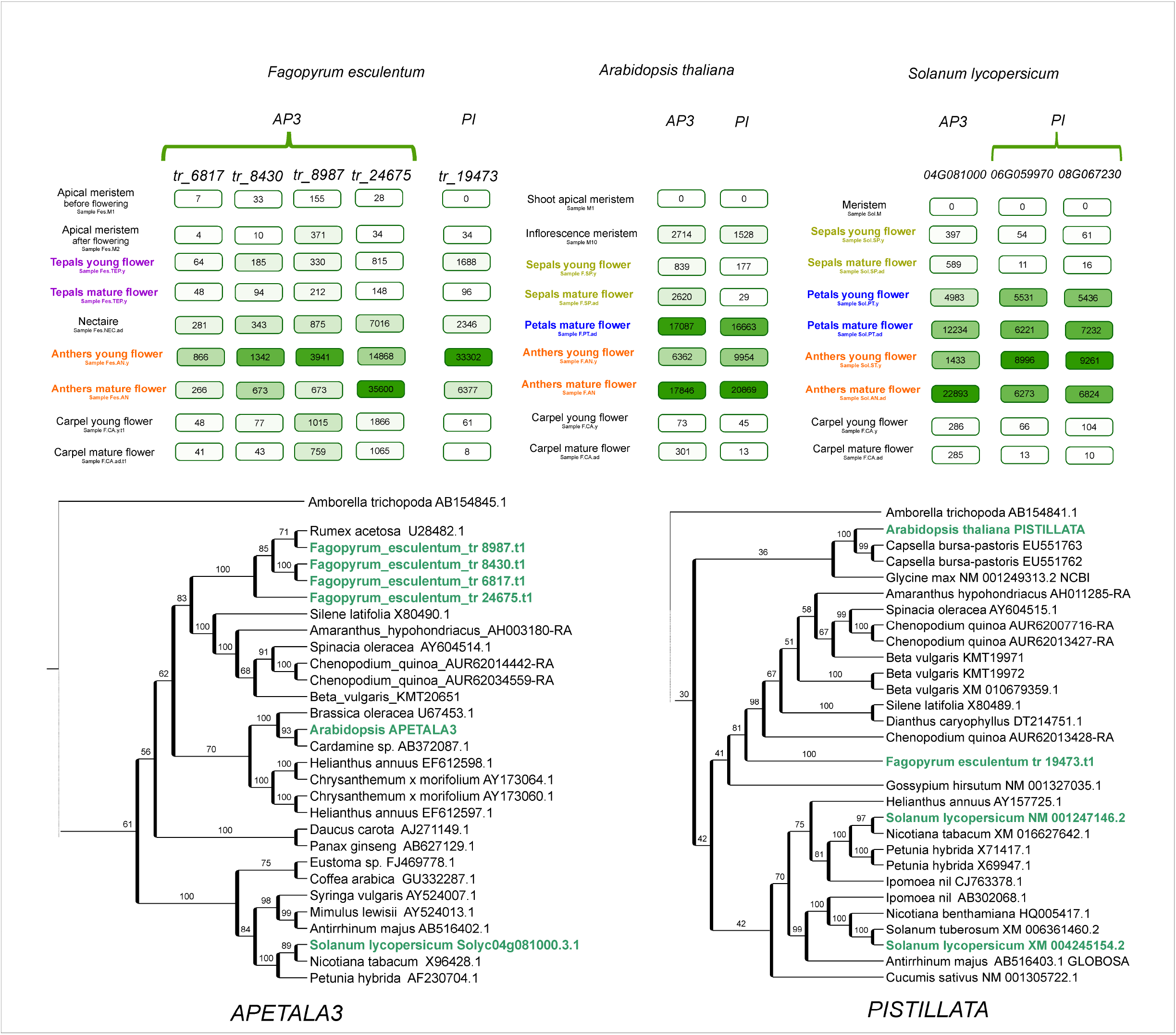
a) Phylogeny and expression patterns of the AP1 orthogroup in *Arabidopsis*, buckwheat and tomato. Numbers in boxes denote normalized read counts. b) Phylogeny and expression patterns of the TIL1-2 orthogroup in *Arabidopsis*, buckwheat and tomato. Numbers in boxes denote normalized read counts.

Another example is provided by the buckwheat tr_14963 gene, which encodes an ~2000 aa protein with high similarity to the DNA polymerase epsilon subunit. It exists in a single copy in most plants, excluding those that recently underwent a whole-genome duplication (Figure 4b). However, in *A. thaliana,* there are two genes, *TIL1* (*AT2G27120*) and *TIL2* (*AT2G27120*), corresponding to this single gene. They show very close similarity values (56%) to the buckwheat orthologue, making it impossible to determine which of the *TIL* genes corresponds functionally to the buckwheat tr_14963 gene. In contrast, the expression profiles show clear differentiation: *TIL1* presents a more similar pattern to the buckwheat gene, where the maximal expression level is associated with growing tissues, while *TIL2* exhibits a narrow pattern and is expressed almost exclusively in the anthers (Figure 4b).

### The transcriptome atlas provides a clue regarding the origins of petaloidy in buckwheat

An important characteristic of flowering plants is the perianth – the organs that surround the reproductive structures of the flower and functions in protection and/or the attraction of pollinators. The perianth can be either double, being differentiated into a calyx (sepals, usually small and green, protect the developing floral organs) and corolla (petals, large and showy, attract pollinators) or undifferentiated, in which a single type of organ resembles either petals or sepals. A double perianth is typical for eudicots (including the model species *A. thaliana*), while an undifferentiated perianth is usually found in monocots. The well-known ABC model of floral organ identity suggests that this structure is controlled by the combinatorial action of three classes of TFs – A, B and C. According to this model, sepal identity is mediated by the A class genes, carpel identity by C class, petal identity by classes A and B and stamen identity by classes B and C (Coen and Meyerowitz, 1991). The undifferentiated petaloid perianth has been hypothesized to be attributable to the expansion of the B class (van Tunen, Arjen *et al.*, 1993); this hypothesis has been experimentally corroborated in several monocot species (Kanno *et al.*, 2003; Nakamura *et al.*, 2005). In Caryophyllales, perianth structure is highly diverse; this group includes species with both double and undifferentiated perianths; the latter type is presumably an ancestral character for the whole order (Brockington *et al.*, 2009). Buckwheat and other species from the family Polygonaceae exhibit an undifferentiated perianth. In *F. esculentum,* the perianth is petal like, white or pink, while in several other buckwheat species, it is green, sepal like. A plausible hypothesis is that in *F. esculentum,* class A and B genes are active in the perianth, while in species with a sepal-like perianth, only class A genes are active. The availability of the genome sequence in combination with a detailed transcriptome map allows this hypothesis to be tested. The search for orthologues of B-class genes showed that the buckwheat genome carries four orthologues of *AP3* and one orthologue of *PI* (Figure 5). We searched for their expression in different floral organs, and we found that B is not expressed or is expressed at background levels in the perianth or only in stamens. In tomato, the expression pattern of B-class genes is compatible with the ABC model; the conservation of the function of *AP3* and *PI* orthologues (the latter is duplicated in tomato) was previously shown in direct RNAi experiments (Guo *et al.*, 2016; de Martino *et al.*, 2006). Tomato belongs to the asterids, while *Arabidopsis* belongs to the rosids. Buckwheat is more closely related to the asterids, which suggests that the control of petal identity by the B-class genes was lost in an ancestor of buckwheat after its divergence from asterids. Earlier, we suggest that B-class genes are not involved in petaloidy in buckwheat based on the phenotype of a buckwheat homeotic mutant (Logacheva *et al.*, 2008); this hypothesis now gains support from the obtained expression profiles. Studies on the complementation of *Arabidopsis ap3* and *pi* mutants via the overexpression of buckwheat *AP3* and *PI* genes also show limited conservation of function (Fang *et al.*, 2015; Fang *et al.*, 2014). An alternative programme of petal development, which is not dependent on B-class genes, was also found in a family of the core Caryophyllales, Aizoaceae (Brockington *et al.*, 2012).

**Figure 5.**
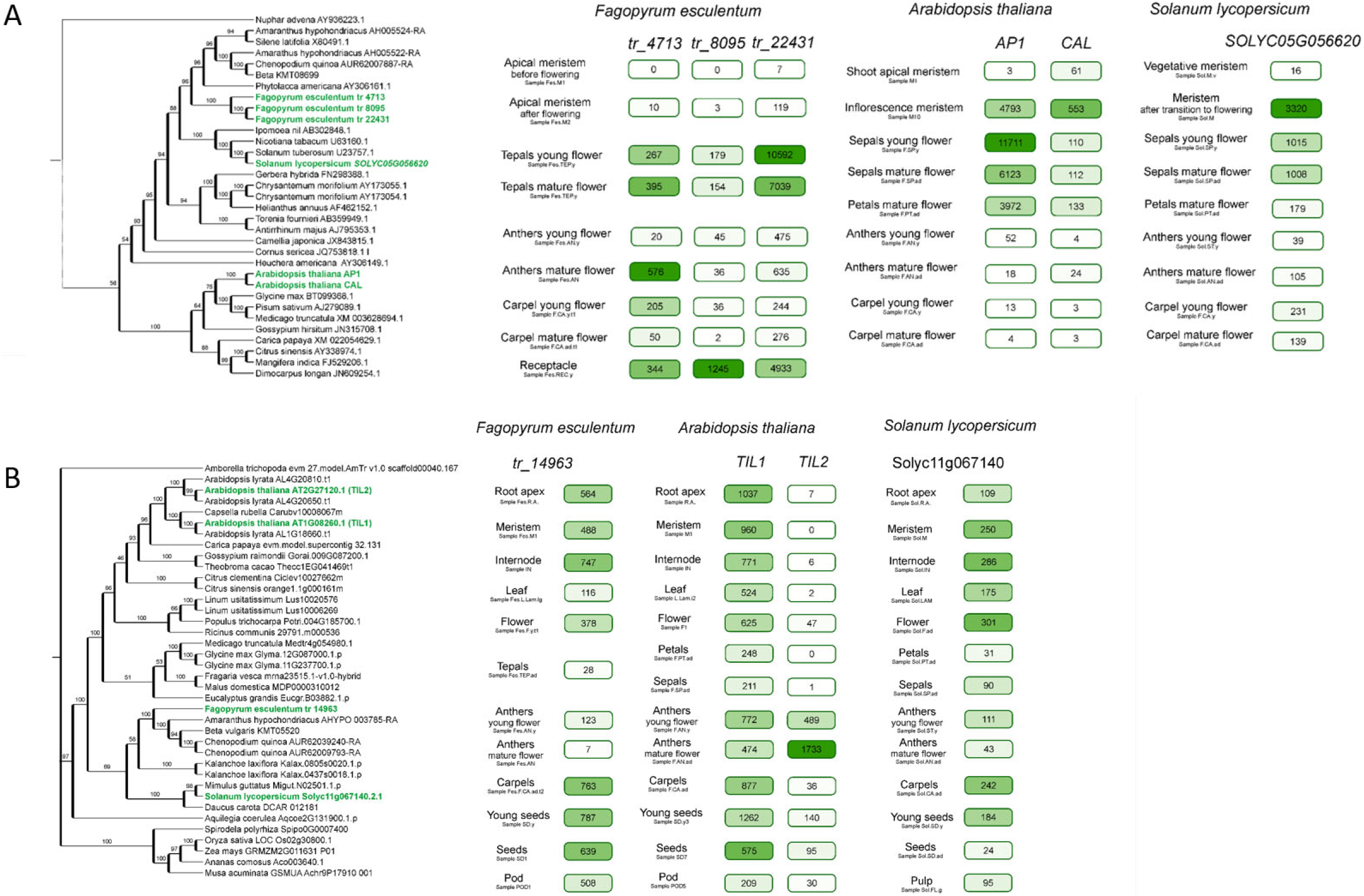
Phylogeny and expression patterns of the orthologues of the B-class genes *APETALA3* and *PISTILLATA* in *Arabidopsis*, buckwheat and tomato, with a focus on floral organs. Numbers in boxes denote normalized read counts.

The most notable features of petaloidy in buckwheat tepals are white colour (while the sepals are usually green) and the shape of epidermal cells, which are conical (Hong *et al.*, 1998), as found in the petals of most plants (Whitney *et al.*, 2011). Studies in the model organism *Antirrhinum majus* have shown that the major role in the development of conical cells is played by the *MIXTA* gene, encoding an R2R3-MYB transcription factor (Glover *et al.*, 1998; Perez-Rodriguez, 2005). Additional factors from the MYB family that are closely related to *MIXTA* control other aspects of this process (Perez-Rodriguez, 2005). *Arabidopsis* does not have a *MIXTA* orthologue, but two genes from a sister clade, *MYB106* (*AT3G01140*) and *MYB16* (*AT5G15310*), as well as several other MYB family TFs, participate in the control of epidermal cell development (Oshima *et al.*, 2013). The evolutionary history of *MIXTA*/*MIXTA-like* genes is complicated, being shaped by multiple duplications and gene losses, particularly in the *MIXTA* clade (Brockington *et al.*, 2013). In the buckwheat genome, we identified 118 genes with significant similarity to MIXTA/MYB16/MYB106 (Table S9). Six of them showed an expression pattern compatible with their role in the determination of conical cells, with prevalent expression in developing tepals (Figure S4). One of these transcripts – tr_18111 – exhibit a protein motif that is characteristic of MIXTA, MYB16 and MYB106. Phylogenetic analysis also indicates that tr_18111 falls into the clade of *MIXTA-like* genes (Figure S5). This strongly supports the hypothesis that petal conical cell identity is controlled by *MIXTA/MIXTA-LIKE* genes in buckwheat as well. Within *Antirrhinum, MIXTA* has been shown to be under the control of B-class genes in *A. majus* (Perez-Rodriguez, 2005; Manchado-Rojo *et al.*, 2012), and this regulatory contour seems to be conserved as far as the monocots (Pan *et al.*, 2014). The absence of B-class gene activity in buckwheat tepals along with the presence of conical cells suggests that this link is not universal. Notably, the morphology of tepal epidermal cells differs within the genus *Fagopyrum*: they are conical in *F. esculentum* and *F. tataricum* (though the latter has green “sepaloid” tepals), whereas they are elongated with sinuated walls in other species (Hong *et al.*, 1998). This genus is thus a good model for the study of the genetic control of petal identity and petal epidermal cell structure. The characterization of the genome and expression profiles of *F. esculentum* enables a comparative transcriptomic study of the genus.

## Conclusions

We characterized the 1.5 Gb genome of *F. esculentum*, providing a reference assembly with high contiguity and an improved representation of protein-coding genes. The genome size of *F. esculentum*, which is three times larger than that of its sister species *F. tataricum*, is a product of a “transposon burst” that occurred 0.5-1 Mya. To provide a framework for the functional and comparative genomics of buckwheat, we constructed a comprehensive transcriptome atlas from 46 tissues, organs and developmental stages. We demonstrated that high-resolution transcriptome maps provide new information that allows us to discern the function of closely related genes and to test biological hypotheses regarding gene function and the conservation of developmental programmes.

## Experimental procedures

### Genome sequencing, assembly and assembly quality check

The Dasha cultivar was chosen as the source material for the construction of the reference genome. DNA was extracted from fresh-frozen young leaves free of visible injuries, necrosis and/or presence of pathogens using the CTAB protocol (Doyle and Doyle, 1987). For Illumina sequencing, a fragment library was prepared using the TruSeq DNA sample preparation kit (Illumina), with 500 ng of DNA as the input and size selection of fragments in the range of 500–600 bp (corresponding to a 380-480 bp insert length) in 2% agarose gels. Sequencing was performed on a HiSeq2500 instrument with HiSeq Rapid 500 cycling reagents at the University of Illinois at Urbana-Champaign (Roy J. Carver Biotechnology Center). Mate-pair libraries were prepared with a Nextera Mate-pair sample preparation kit (Illumina) with 5000 ng of DNA as the input. Size selection was performed in agarose gels; three lengths were selected: 3–4 kbp, 5–7 kbp and 8–10 kbp. Mate-pair libraries were sequenced on a HiSeq2000 instrument with a TruSeq SBS 200 cycle kit. For sequencing on the Pacific Bioscience platform, DNA was extracted in the same way indicated above and sequenced with a SMRT cell on a Sequel II instrument with CCS read settings at the DNALink facility.

The assembly process consisted of three stages: contig assembly, scaffolding and gap closing. For contig assembly, a Newbler assembler was used. Newbler v. 2.9 was run on a 4 Intel(R) Xeon(R) CPU E7-4830 v2 2.20 GHz computer with 2 TB RAM with the following parameters: “-large -ml 150 -mi 85 -het”. These parameters specified a minimal overlap length of not less than 150 bp and minimal overlap identity of 85%. The “long” parameter indicates a large genome (>100 Mb), and the “het” parameter indicates heterozygosity. The resulting contigs with lengths greater than 1000 bp were retained for scaffolding. Scaffolding was performed using Platanus v. 1.0.0 (Kajitani *et al.*, 2014) (subprogramme scaffold) with the parameter “−l 3”, indicating that contig joining should be supported by at least 3 mate-pair links. The closing of the gaps was carried out in two stages. In the first stage, gaps were closed with Platanus version 1.0.0 software (with default parameters) and Illumina mate-pair data. In the second stage, gaps were closed with PacBio CCS data and LR_Gapcloser software (Xu *et al.*, 2019) with the parameter “-r 5”.

### RNA extraction and sequencing

For RNA-seq analysis, RNA was extracted from a set of different organs and developmental stages using the RNEasy Plant Mini kit (Qiagen, Netherlands) with the modifications described previously (Logacheva *et al.*, 2011). RNA quality was checked using a Bioanalyzer2100 (Agilent, USA); samples with RIN > 7 were employed for subsequent analysis. For Oxford Nanopore Technologies sequencing, RNA was converted to cDNA using a Mint cDNA synthesis kit (Evrogen, Russia). The primers for reverse transcription included custom barcodes unique to each sample; thus, the samples corresponding to different organs/stages were pooled after reverse transcription. cDNA was amplified via 24 cycles of amplification. Amplified cDNA was used as the input for library preparation via a standard protocol for genomic DNA with an LSK-309 kit. Sequencing was performed on a MinION system with a 9.5.1 flow cell, and base calling was performed with Guppy. For Illumina sequencing, the libraries were prepared using a TruSeq RNA sample preparation kit (Illumina, USA) following the manufacturer’s protocol and sequenced on a HiSeq2000 instrument (Illumina, USA) with 50-bp single reads.

### Annotation

The assembled scaffolds after the first stage of the gap-closing procedure for *F. esculentum* were repeat masked with *RepeatMasker* (ver. 3.3.0) before annotation (http://www.repeatmasker.org). Gene models that overlapped with any of the repeat regions were given a score penalty. RNA-seq data were mapped with *STAR* (ver. 2.4) on the masked genome assembly, and the obtained alignment files were used to generate hints for *Augustus* consisting of hints about the intron start and end sites and transcribed regions. The initial set of genes was predicted ab initio by running the *Augustus* (ver. 3.0.1) gene prediction program with different parameters (species model: *Arabidopsis*/tomato; hints: with/without hints, 4 runs in total) and *GeneMark*. ES (ver. 2.3). All predicted gene models were combined. All predicted gene models were evaluated and ranked according to the following criteria: 1) a small positive bonus was added to the gene model if the intron or exon was supported by RNA-seq data; 2) a positive bonus proportional to the quality and significance of *blastp* alignment with known proteins from the following datasets: *A. thaliana* (maximal bonus if the alignment was significant), SwissProt (smaller bonus) and the NCBI non-redundant database (smallest bonus), was added to the gene model; 3) large negative penalties were applied if the model intersected with repeat regions or showed a hit with a TE. Models were sorted by score. The best-scoring models were picked first, and only one best-scoring isoform was included in the final annotation (no overlapping models). Orthogroups were obtained using Orthofinder software version 2.4.0 (Emms and Kelly, 2019). The annotation of TFs was performed using PlantRegMap (Tian *et al.*, 2019). The completeness of the assembly was estimated using BUSCO version 4.0.6 software (Simão *et al.*, 2015) for the eudicot lineage and the Viridiplantae lineage.

### Transposable element analysis

To compare the repeatome composition between species, 4 000 000 *F. esculentum* and 1 600 000 *F. tataricum* high-quality paired-end Illumina genomic reads were randomly sampled, and suffixes and prefixes were added using a Python script (https://github.com/Kirovez/RepeatExplorer_scripts/blob/master/prepareReadsREv2.py). The reads were combined into a single FASTA file and used for RepeatExplorer (Novak *et al.*, 2013) and TAREAN analysis (Novák *et al.*, 2017). The locally installed version of RepeatExplorer containing the built-in TAREAN program was run with the following settings: -p -c 150 -C -r 400000000 -P 2. The automatic annotation data were used for repeat classification. Clusters corresponding to organelle DNA were removed, and the portion of the genome consisting of repeats was corrected according to the new number of reads. The table containing the information about cluster annotation and read number was analysed in Rstudio Version 1.2.1335 (http://www.rstudio.com/) with R version 3.6.0 using custom scripts with relevant packages, including ggplot2 (Wickham, 2009) and data.table (https://cran.r-project.org/web/packages/data.table/index.html). For the comparison of the repeatome composition between FE cultivars, 2 000 000 high-quality paired-end Illumina genomic reads were randomly selected, and RepeatExplorer was run as described above. A heatmap was constructed using the ComplexHeatmap R package (Gu *et al.*, 2016).

For the genome-based analysis of LTR RTEs in the two species, the genome assembly of *F. tataricum* (http://mbkbase.org/Pinku1/FtChromosomeV2.fasta.gz, (Zhang *et al.*, 2017)) was downloaded. The LTRharvest tool with default parameters (Ellinghaus *et al.*, 2008) was used to predict putative coordinates of LTR RTEs in the genomes. The obtained gff3 file was sorted and used as the input for the LTRdigest program (Steinbiss *et al.*, 2009) with the following settings: - aaout yes -pptlen 10 30 -pbsoffset 0 3 -pdomevalcutoff 0.001. Hmm repeat profiles of RTE domains were downloaded from the GyDB database (Llorens *et al.*, 2011). The gff3 file from LTRdigest analysis was parsed using a custom python script (https://github.com/Kirovez/LTR-RTE-analysis/blob/master/LtrDiParser_v2.2.py) to extract the sequences of full-length LTR RTEs (those possessing similarity to GAG, RT, RH, AP and INT domains) and their LTRs. TEsorter software (Zhang *et al.*, 2019) was used for RTE classification. The insertion time was calculated with the formula T = k/2r, where k is the distance between LTRs estimated via Kimura’s two-parameter method (Kimura, 1980) and r is the mutation rate. A mutation rate of 1.3 × 10−8 substitutions per site per year (Ma and Bennetzen, 2004) was used. Parameter K was calculated as 0.5 log((1 - 2p -q) * sqrt(1 - 2q)), where p is the transition frequency and q is the transversion frequency. The transition and transversion frequencies were estimated after the alignment of 5’ and 3’ LTR sequences with clustalw2 software (http://www.clustal.org/clustal2). The insertion time analysis was automated with a custom Python script (https://github.com/Kirovez/LTR-RTE-analysis/blob/master/TEinsertionEstimator.py).

### Gene expression analysis

The total number of reads mapped on a given gene (TGR) was used as a measure of the expression level. The “DESeq” (Anders and Huber, 2010) median approach was used for the normalization of gene expression levels according to library size. We used a threshold of 5 or a more normalized TGR in both biological replicates to identify the expressed genes. “DESeq2” (Love *et al.*, 2014) was used for differential expression analysis with the following settings: a false discovery rate (FDR) <0.05 and a fold change ≥ 2. The differential expression (DE) score was defined as reported previously (Klepikova *et al.*, 2016). Gene expression pattern width was assessed as described by Penin *et al.* (2019).

### Phylogenetic analysis

Nucleotide sequences were used for phylogenetic analysis. Alignment was performed using MUSCLE (Edgar, 2004), and further was conducted processing using GBlocks with “less stringent” settings (Castresana, 2000). For phylogenetic tree reconstruction, we used IQ-tree (Minh *et al.*, 2020) with the GTR model and 1000 bootstrap replicates.

## Supporting information

Data Sheet 1

Data Sheet 2

## Data availability statement

Genome assembly and raw sequencing reads for the Dasha cultivar (reference genome) are available in the NCBI database under Bioproject # PRJNA487881. Raw reads from buckwheat cultivars and *F. esculentum* ssp. *ancestrale* are available under Bioproject # PRJNA627307. The transcriptome reads are available from NCBI SRA, and their accession numbers are listed in Table S5.

## Acknowledgments

The study was supported by Russian Foundation for Basic Research, project # 18-29-13017 (development of a transcriptome map) and Russian Science Foundation, project # 18-76-10018. We are grateful to Dr. Alexey Kondrashov for the access to sequencing and computational facilities.

## Author contributions

AAP constructed transcriptome libraries and coordinated the part of the work on the transcriptome analysis, ASK performed genome assembly, AVK performed gene expression analysis, IVK analysed the repeatome and participated in writing, ESG performed genome annotation, ANF provided plant material, MDL conceived and coordinated the study, constructed genome libraries, performed phylogenetic analysis and drafted the manuscript.

## Conflict of interest

The authors declare that they do not any conflict of interest.

## Supporting figures

Figure S1.

Repeat content in *F. esculentum* spp. *ancestrale* and different buckwheat cultivars. DAS – Dasha, BAS – Bashkirskya krasnostebelnaya, DEM – Demetra, DEV – Devyatka, DIA – Dialog, DIZ – Dizajn, KAR – Karadag, KAZ – Kazanka, KUJ – Kujbyshevskaya, DRR/YAS – the sample from the study by Yasui et al. 2016, cultivar unknown, presumably of Japanese origin, KIT/SHI – Shinanonatsusoba, FEA - *F. esculentum* spp. *ancestrale*.

Figure S2

The distribution of Shannon entropy across expressed genes of *F. esculentum.*

Figure S3.

Heatmap showing the expression profiles of the genes – orthologues of the stably expressed genes in *Arabidopsis thaliana*.

Figure S4

Heatmap showing the expression of buckwheat MYB genes sharing significant similarity with MIXTA/MYB16/MYB106.

Figure S5

Phylogenetic tree of angiosperm MIXTA/MIXTA-like genes.

## Supporting tables

Table S1

Data generated for the de novo assembly of the buckwheat genome and accession numbers of raw reads

Table S2

Statistics of the assembly

Table S3

Statistics for the presence of universal single-copy orthologues for the earlier assembly of the buckwheat genome and for the assembly reported in our study

Table S4

Statistics of the protein-coding gene annotation

Table S5

Transcription factor genes annotated in the buckwheat genome

Table S6

List of the samples included in the transcriptome mapping procedure and accession numbers of the raw reads

Table S7

Samples included in the transcriptome mapping procedure and correlations

Table S8

GO enrichment of the genes with broadest and narrowest expression patterns

Table S9

Buckwheat MYB genes selected based on similarity to MIXTA and MYB16/MYB106

## Notes

### Competing Interest Statement

The authors have declared no competing interest.

