## Supplementary figures and images for "High-resolution transcriptome atlas and improved genome assembly of common buckwheat, *Fagopyrum esculentum*"

### Data Sheet 1

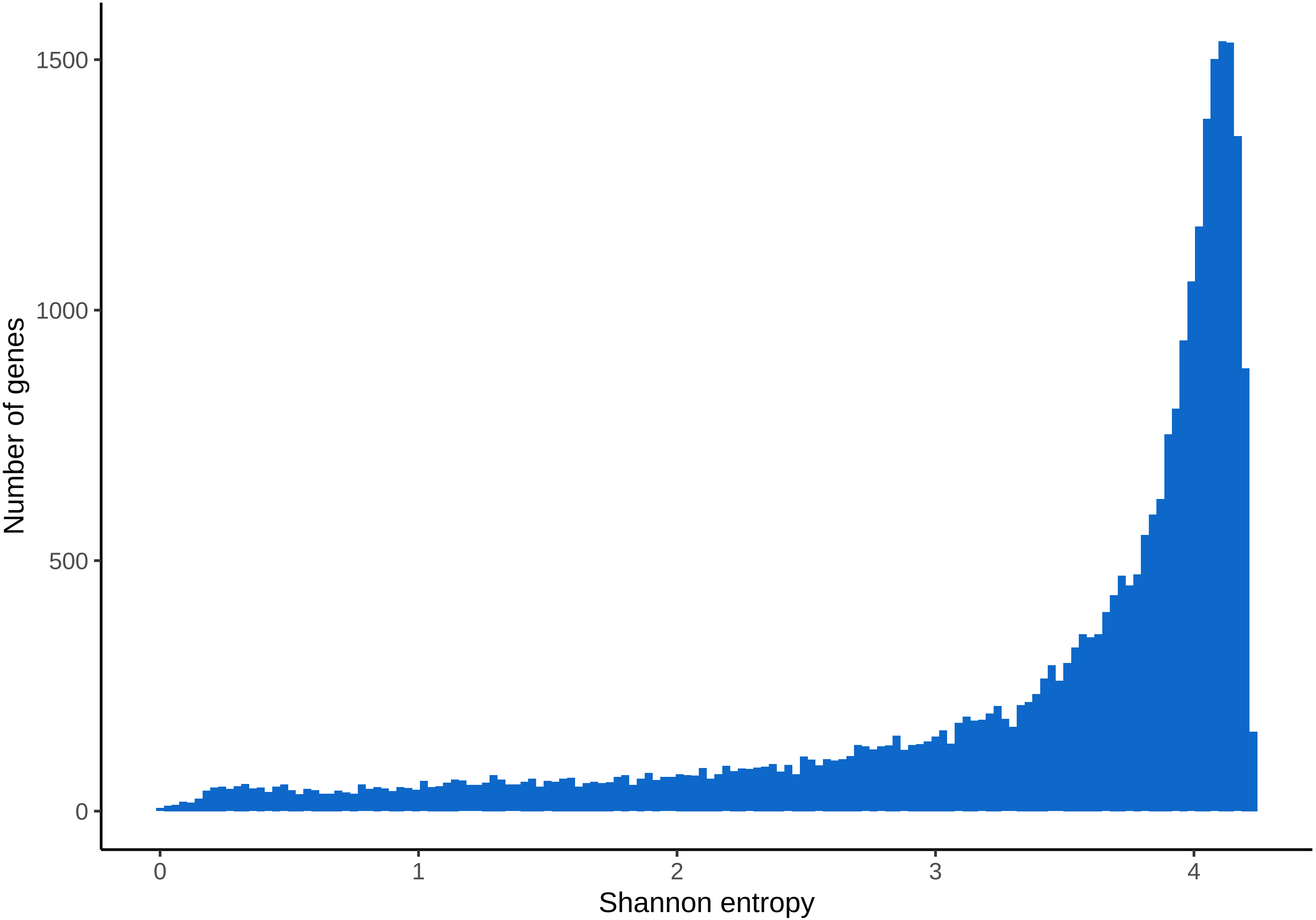

### Data Sheet 2

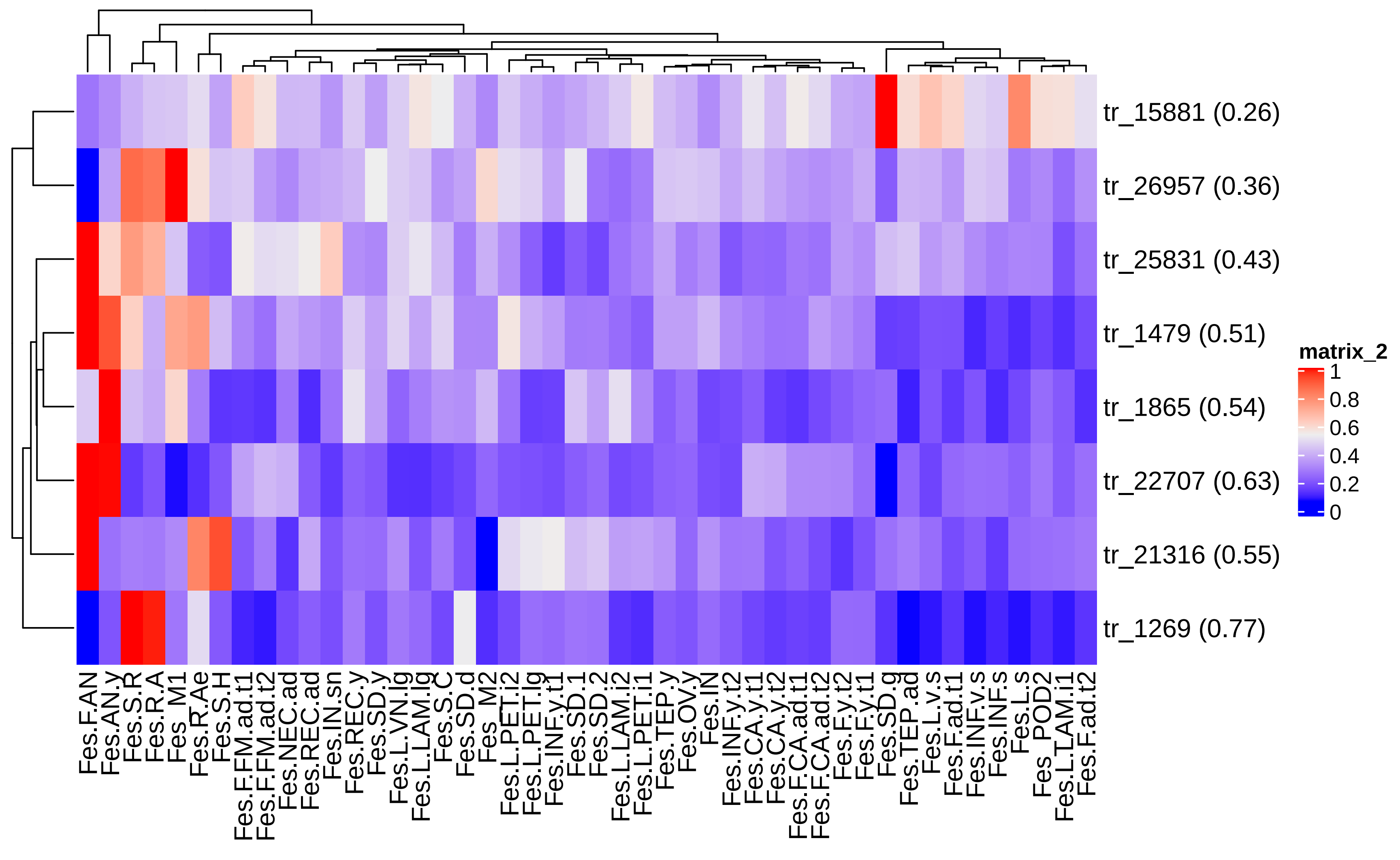

### Figure S1

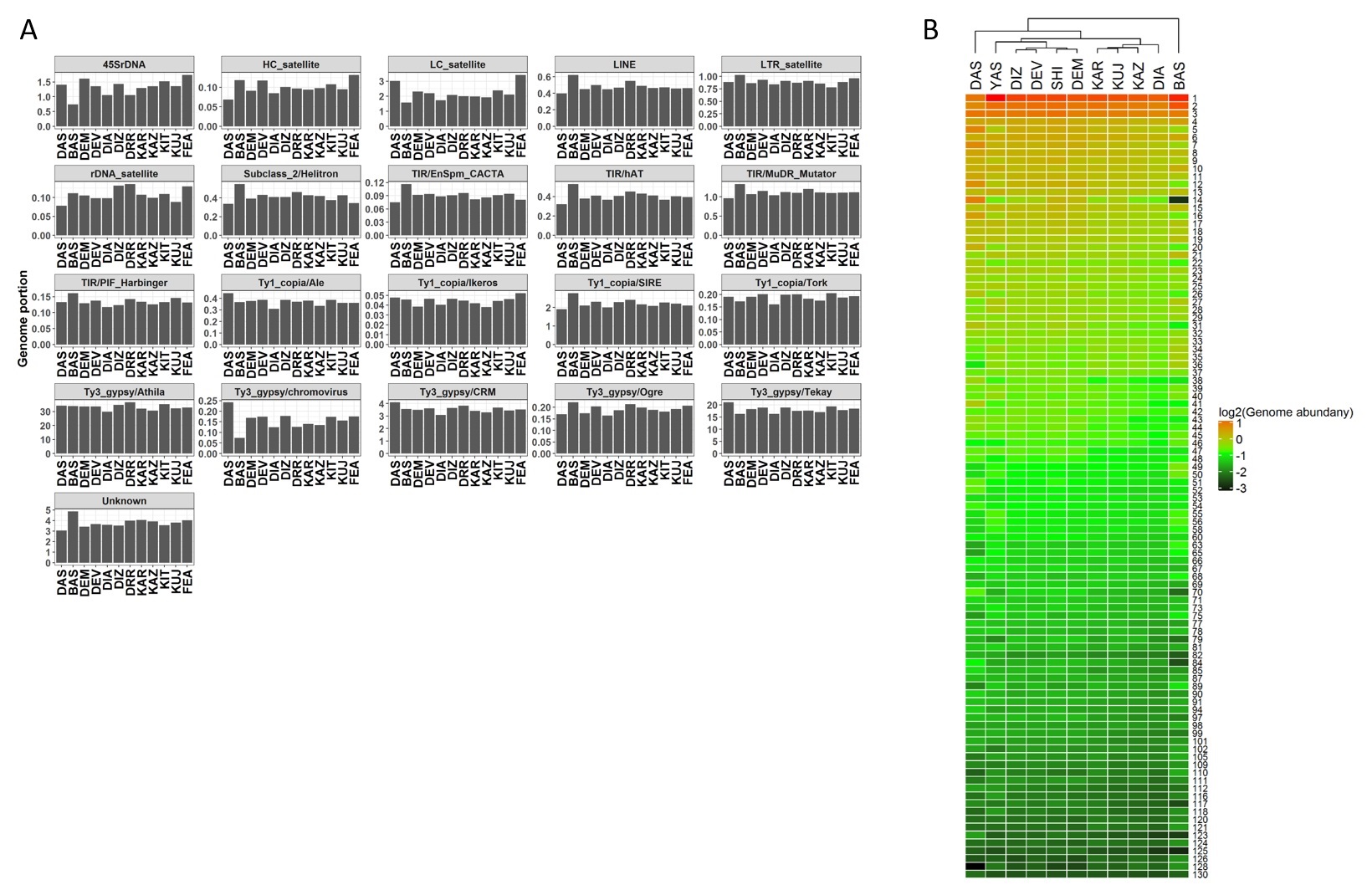

### Figure S2

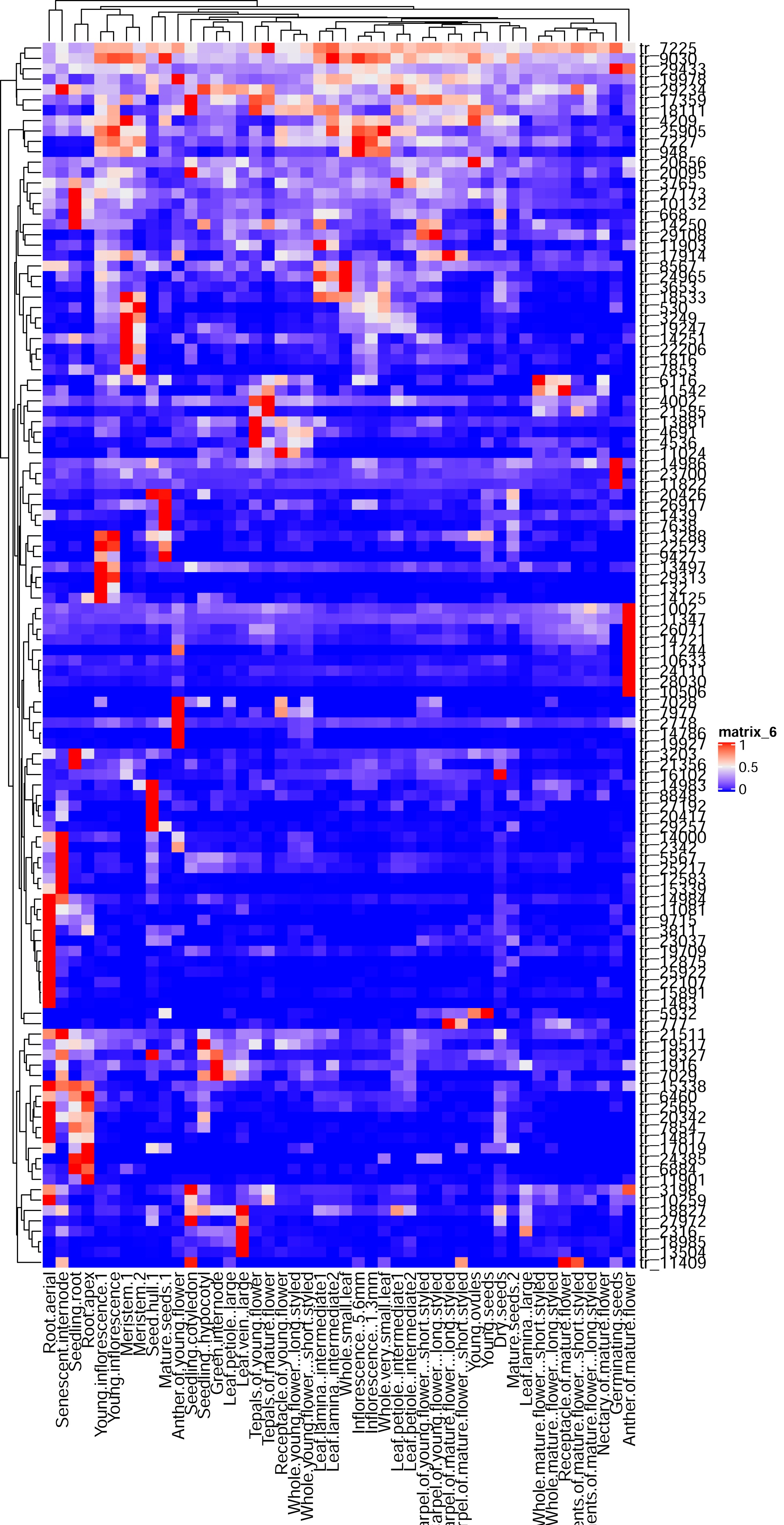

### Figure S3

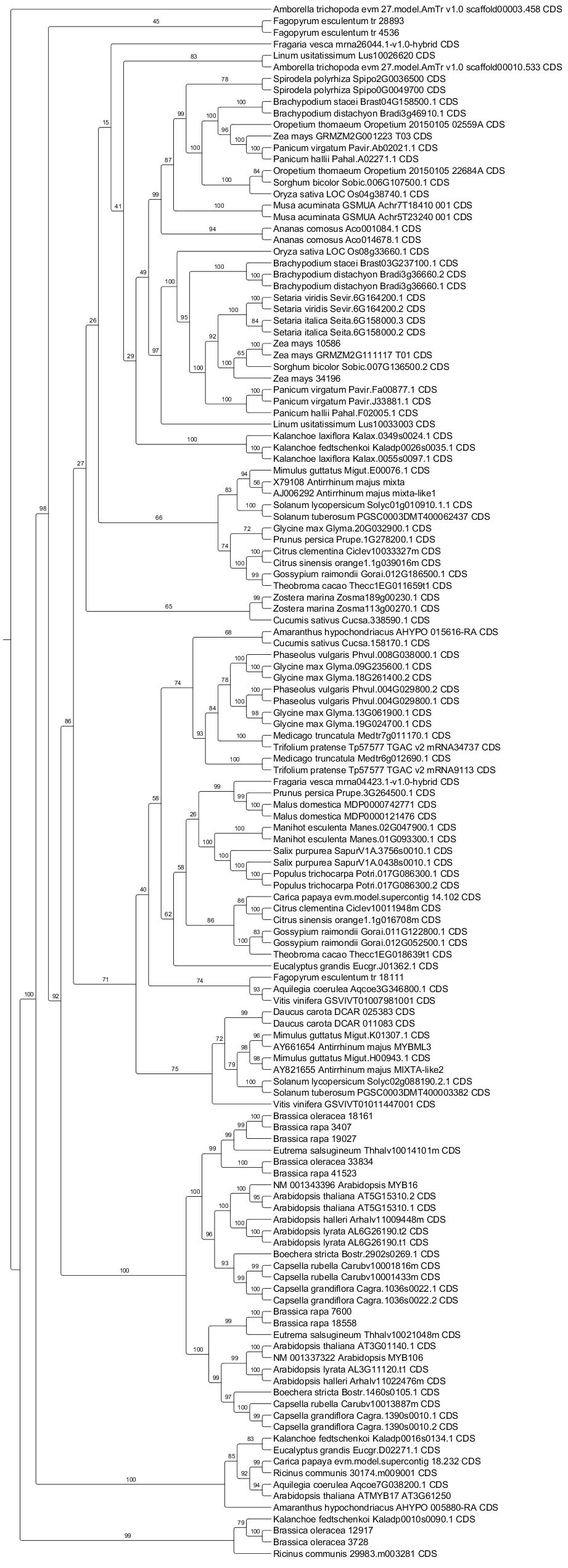
